# Identification of Microbial Flora and Isolation of Functional Strains Related to the Degradation of Domestic Waste

**DOI:** 10.1101/2022.03.20.485016

**Authors:** Tianyi Huang, Xinyi Liu, Yeqing Zong

**Affiliations:** Shanghai Starriver Bilingual School, No.2588, Jindu Road, Shanghai, China; Bluepha Co., Ltd., Bldg.4-F5-C501, NIU ZONE, 430 Linqing Rd, Yangpu Dist, Shanghai 310110, China

**Keywords:** Household food waste, Compost, Microbiota, Functional strains

## Abstract

A large amount of food waste is currently generated due to rapid global population growth, which results in environmental pollution and economical burdens for waste disposal. Composting is one of the most environmentally friendly and cost-effective methods for food waste treatment. This study aims to give insight into microbial community structure and functional strains of household food waste compost, which might optimize the composting process and help increase food waste treatment efficiency. The microbial community is characterized via 16S rRNA sequencing. The dominant microbes are assigned to the three phyla of *Firmicutes*, *Proteobacteria* and *Bacteroidetes*, and the main fungi are assigned to the phylum of *Ascomycota*. We isolated some functional microbes from the compost microbiota, including *Alcaligenes faecalis*, *Paenochrobactrum glaciei*, *Lysinibacillus fusiformis*, and *Proteus terrae*. Moreover, we detected lipase and cellulase activities in the compost liquid. However, the measured lipase and cellulase activity is low, and further enhancement of the lipase activity in the food waste compost might increase waste treatment ability. Therefore, we isolated a cellulase and lipase harboring *Bacillus cereus* strain from municipal a wastewater treatment plant. The *B. cereus* strain have high-level lipase activity, which could be used for food waste treatment in the future.

## 1. Introduction

Along with the rapid population growth, large amounts of food waste are generated. It is estimated that 1.3 billion tons of food waste are produced each year (United Nations, 2018). The untreated food waste leads to environment pollution (Zhao et al., 2017). However, food waste disposal creates economical burdens for society. Take Shanghai as an example, the expenditure spent on food waste transformation and treatment reach as high as 380 yuan/ton. Therefore, developing eco-friendly and low-cost food waste treatment strategy is of great interest.

Currently, four main methods, composting, landfill, anerobic digestion, and thermal conversion, are used for food waste treatment. Composting requires the addition of moisture during processing and produces odors (Cerda et al., 2017). The final product of composting can be used as fertilizer, and fewer green-house gas emissions and leachate are generated (Adhikari et al., 2009). Landfill can dispose of food waste efficiently (Kibler et al., 2018), but it generates leachate which can contaminate surface and ground water (Kibler et al., 2018), and the biogas production efficiency is low (Thyberg et al., 2017). Anaerobic digestion biosynthesizes methane (Kibler et al., 2018), but it often performs at low-loading rates due to system instability (Xu et al., 2017). Thermal conversion treatment of food waste can be usedto generate electricity (Kibler et al., 2018), but it produces a high carbon-footprint (Jeswani et al., 2013).

The most preferred food waste treatment strategywould reduce or avoide waste during treatment. Composting is an efficient food waste valorization method, which degrades organic matter under aerobic and anaerobic processes. The final composting product is rich in inorganic nutrients, which can be used as plant fertilizer. Therefore, there is great value in improving efficiency to the composting process. The contents in the food waste are highly variable and heterogeneous (Cerda et al., 2017), and the treatment of food waste are often conducted by large food companies. Moreover, current studies in composting mainly focus on improving the efficiency of the composting process using thermophilic strains (Antunes et al., 2016). However, household food waste composting and its microbial compositions targeted on organic matter degradation have not been widely evaluated.

The purpose of this research is to recover microbiota and functional strains involved in the composting process with designed food waste as substrate. This research might help improve composting efficiency within the household environment, which could potentially lower the costs for food waste treatment. Our research purposes and goals are listed as follow,

1. Complete the composting process with designed food waste.
2. Identify and isolate functional strains from the compost.
3. Analyze the microbial structure of the compost samples.
4. Characterize the cellulase and lipase activities, and try to isolate functional strains for future food waste composting.

## 2. Materials and Methods

### 2.1 Composting substrate

Three composting boxes were used for the composting process each imitating a different scenario of food waste, and the three composting boxes were named as FW-B1, FW-B2, and FW-B3, respectively. The food waste compositions of these three boxes were designed according to a previous study (Song et al., 2015). FW-B1 and FW-B3 had the same composition, but commercial composting strains were added to FW-B1. The commercial composting strains were provided by EMRO (http://www.emrojapan.com), which harbor lactic acid bacteria, yeast, and phototrophic bacteria. The vegetables, meal, meats & fish, fruit and eggshell each composed 52.2%, 22.2%, 12%, 6.1% and 2% in FW-B1 and FW-B3. The FW-B2 contained 0.05 kg rice, 0.08 kg oil crops, 0.68 kg vegetables, 0.14 kg fruit, 0.2 kg meal, 0.03 kg meats & fish, and 0.003 kg other substances. The FW boxes were kept at room temperature, and the composting lasted for 81 days.

### 2.2 16S rRNA and ITS sequencing

The 16S rRNA gene is about 1.5 kb in length, and it contains 10 constant regions and 9 variable regions, which can be used as a marker gene for prokaryotes. The ITS rDNA genes are unique to fungi, which codes for the small subunit of fungal ribosomes. The V4 region of 16S rRNA gene was sequenced, and the primer pair of 515F 5’-GTGCCAGCMGCCGCGGTAA-3’and 806R 5’-GGACTACHVGGGTWTCTAAT-3’was used. The raw sequences were processed, and the obtained sequences with more than 97% similarity were classified as one operational taxonomic unit (OTU) (Wei et al. 2020). The microbial structures of the three microbial communities, and the alpha diversity of these three composting samples were calculated. The obtained 16S rRNA high-throughput sequencing data of FW-B1, FW-B2 and FW-B3 were deposited in SRA database with the accession numbers of SAMN21356052, SAMN21356053 and SAMN21356054 respectively.

### 2.3 Compost strains isolation and cultivation

1 mL liquid samples were collected from each box and inoculated into 250 ml Erlenmeyer flasks with 90 ml sterilized water. The samples were cultivated at 30 °C and 160 r/min for 30 minutes. The culture was diluted to 10^-1^ to 10^-6^ with sterilized water. Each dilution was spread on LB solid culture media consisting of 10 g tryptone, 5 g yeast extract, 10 g NaCl, 20 g agar (Zhao et al., 2017). Distinct colonies were marked and recorded, then they were transferred to fresh LB culture media for further streaking. The process was repeated for two times, and the DNA of the isolated colonies were extracted and 16S rRNA gene were sequenced. All the obtained 16S rRNA genes were deposited in Genbank database with the accession numbers of MZ960169—MZ960179. The same procedure was used to isolate functional strains from municipal wastewater treatment microbiota, and the 16S rRNA gene accession number of the isolated strain was deposited in Genbank database with the accession number of OK058270.

### 2.4 Methods for testing starch, lipid, and cellulose degradation ability

Verification of organic matter degradation ability was conducted via cultivation of the isolated strains on the medium with cellulose, starch, or lipid as substrate. The cellulose medium (1,000 ml) composed with 0.05% of K_2_HPO_4_, 0.188% microcrystalline cellulose, 0.025% MgSO_4_, 0.2% gelatin, 0.02% Congo red, and 1.4% agar. The starch medium (1,000 mL) was consisted of 2% of soluble starch, 0.5% NaCl, 0.5% peptone, and 2% agar. The lipid medium (1,000 mL) was consisted of 1% peptone, 1% NaCl, 0.01% CaCl_2_·H_2_O, 1% Tween-80, and 2% agar, and the pH was adjusted to 7.4–7.8. Clearance zones was used as the indicators of degradation activity of each strain (Zhao et al., 2017).

### 2.5 Lipase activity measurement

The medium used for lipase activity measurement is composed with 0.01% MgSO4, 0.1% KH_2_PO_4_, 1% Tween-80, 0.2% glucose, and 0.5% peptone. The isolated strains were inoculated into the medium, and 5% of overnight cultures was inoculated into the cultivation medium. The culture was cultivated at 37 °C for 24 hours at 150 rpm. The obtained culture was centrifuged at 8000 rpm for 15 min, and the supernatant was collected for lipase activity measurement. Tween 20, Tween 60, or Tween 80 were used as the lipase activity testing substrate. The enzyme measurement reaction mixture (3 ml) was consisted with 2.3 ml 50 mM Tris hydrochloride, 0.1 ml 10% of Tween 20/60/80 dissolved in Tris buffer, 0.1 ml of 1M CaCl_2_, and 0.5 ml fermentation supernatant. The protein concentration in the supernatant was measured with the BCA protein assay kit (Sangon Biotech).

During enzymatic activity measurement, 0.5 ml heat-treated samples (95 °C for 10 mins) was used as control samples. The control and reaction mixtures were incubated in a 37 °C water bath for 2 hours. Lipase activity was determined with a spectrophotometer at 400 nm. One unit of enzyme activity (U) is defined as a 0.01 increase in OD after the incubation of the reaction mixture with the fermentation liquid. Lipase activity is expressed as unit of enzyme activity per mg of the enzyme (U/mg) (Serikovna et al., 2013).

## 3. Results

### 3.1 Characterization of Composting process

During the 81-day composting process, temperatures of the compost boxes were monitored. And the composting temperature is room temperature at the range of 18°C-26°C (Figure 1a). The malodorous gas NH_3_ is also monitored during the composting process using a universal pH indicator. If the wetted pH test paper indicates that the leachate of compost is alkaline, it is assumed that the malodorous gas produced by the composts is ammonia, which is alkaline. It is proposed that NH3 was detected throughout the composting process (Figure 1b).

**Figure 1.**
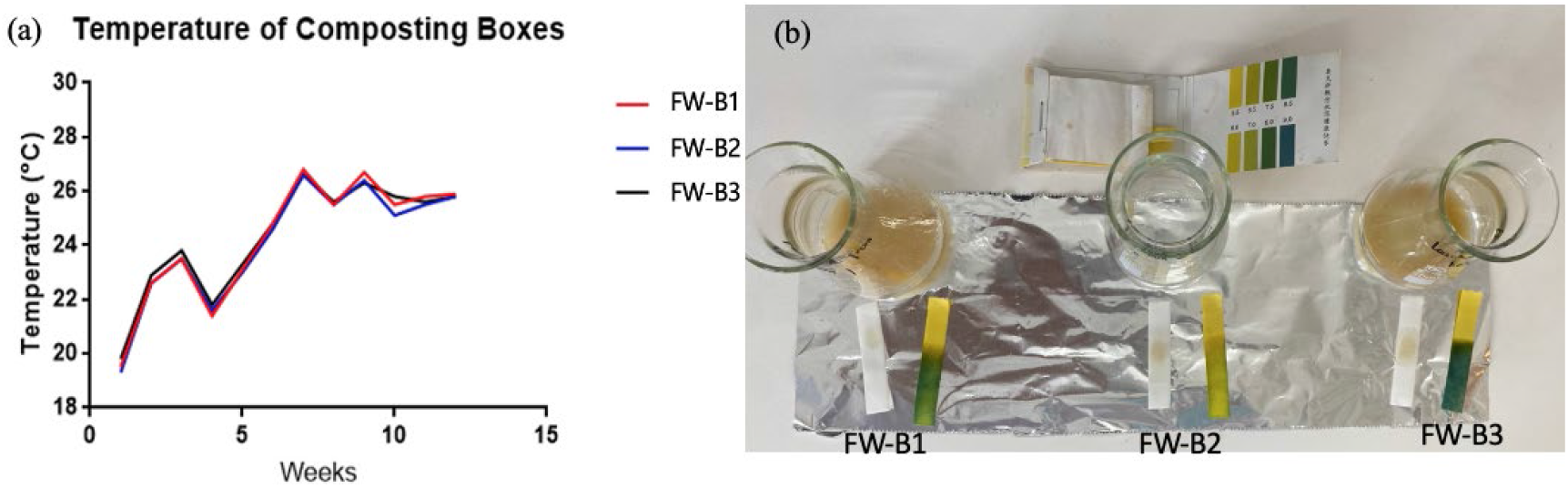
Characterization of composting process. (a) Temperature during composting; (b) Detection of NH_3_ by pH universal indicator

### 3.2 Microbial profiles of the composts

A total of 130,627, 88,249, and 93,319 sequences were obtained from FW-B1 FW-B2, and FW-B3, respectively. Based on the 97% identity classification principle, a total of 473 OTUs were obtained. The composts of FW-B1, FW-B2, and FW-B3 harbored 387, 405, and 423 OTUs, respectively. FW-B1 and FW-B2 have 332 common OTUs; FW-B2 and FW-B3 have 382 common OTUS; FW-B1 and FW-B3 have 344 common OTUs. In fact, most OTUs were common in the three composts, which consisted of 66.6% of the OTUs (Figure 2).

**Figure 2.**
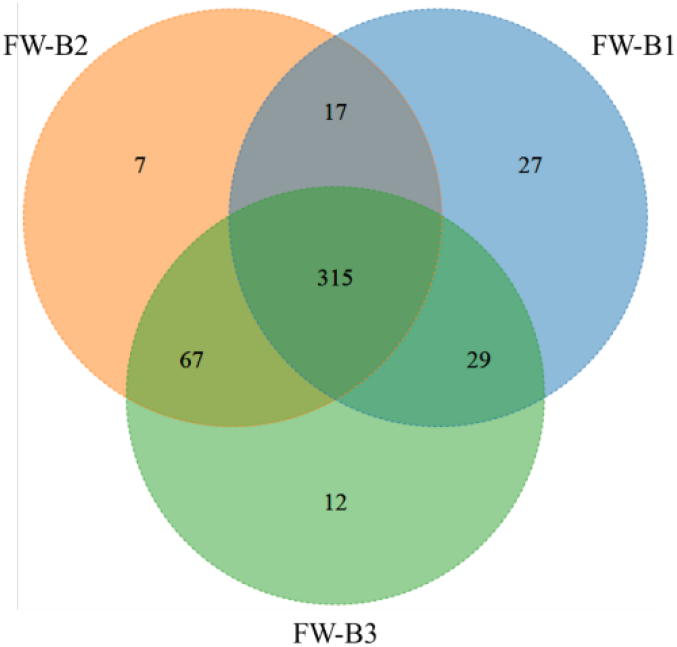
OTU numbers in the three composts.

### 3.3 Microbial distribution at phylum and genus level

For FW-B1, the most dominant phyla were *Proteobacteria* and *Firmicutes*, which composed 47.0% and 42.6% of the total sequences, respectively. For FW-B2, the most dominant phyla were *Firmicutes* and *Proteobacteria*, and they composed 51.9% and 34.2 %, respectively. For FW-B3, the most dominant phyla were *Proteobacteria* and *Firmicutes*, which were same with FB-B1. The compositions of *Proteobacteria* and *Firmicutes* were 58.3% and 20.3%, respectively. Besides these two dominant phyla in these three composts, the third most dominant phylum was *Bacteroidetes*, which composed 10.3%, 12.4%, 21.2% of FW-B1, 2, and 3, respectively. These three dominant phyla composed 99.9% of FW-B1, 98.5% of FW-B2, and 99.8% of FW-B3 (Figure 3a).

**Figure 3.**
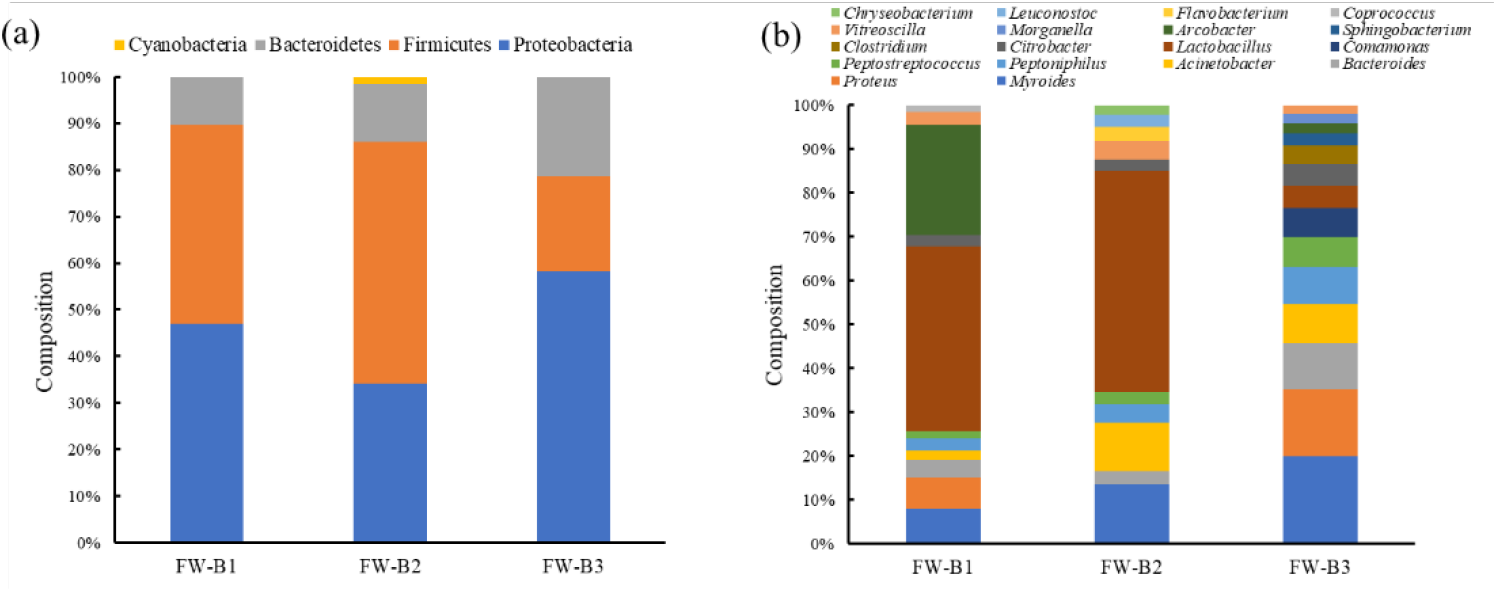
The most dominant microbes of the three microbial communities at (a) phylum and (b) genus level.

At the genus level, the most dominant genus of FW-B1 and FW-B2 was *Lactobacillus*, and the compositions of *Lactobacillus* in FW-B1 and FW-B2 were 32.6% and 25.3%, respectively. The most dominant genus of FW-B3 was *Myroides*, which composed 10.7% of the microbiota. *Myroides* is one of the three most abundant genera in FW-B1 and FW-B2, and it composed 6.8% in FW-B1 and 6.2% in FW-B2. The compositions of the dominant genera in these three microbial communities are different (Figure 3b). In general, the most dominant genera of FW-B1 were *Lactobacillus* (32.56%), *Arcobacter* (19.35%), *Myroides* (6.23%), *Proteus* (5.46%), *Bacteroides* (3.09%), and *Vitreoscilla* (2.40%) (Figure 3b). The most dominant genera of FW-B2 were *Lactobacillus* (25.31%), *Myroides* (6.81%), *Acinetobacter* (5.59%), *Vitreoscilla* (2.20%), and *Peptoniphilus* (2.16%) (Figure 3b). The most dominant genera of FW-B3 were *Myroides* (10.66%), *Proteus* (8.20%), *Bacteroides* (5.59%), *Acinetobacter* (4.87%), and *Peptoniphilus* (4.50%) (Figure 3b).

The alpha-diversity index was also calculated for the bacterial composition from the compost. The alpha-diversity index reflects microbial richness and diversity. FW-B3 have more species than the other two composts (Table 1). FW-B1 have the lowest diversity in these three composts based on the Shannon and Simpson indices. Though the OTU counts were lower than the of Chao1 counts, the Good’s coverage is 100%, indicating that all the microbes in these three composts were covered during high throughput sequencing (Table 1).

**Table 1.** Alpha-diversity indices of the three composts

|  | Richness | Chao1 | Shannon | Simpson | Good's coverage |
| --- | --- | --- | --- | --- | --- |
| **FW-B1** | 388 | 414 | 4.89 | 0.92 | 100% |
| **FW-B2** | 406 | 409 | 6.15 | 0.97 | 100% |
| **FW-B3** | 423 | 428 | 6.34 | 0.977 | 100% |

#### 3.3.1 Cultured Strains Identification

After morphological identification of the bacterial colonies isolated from the three composts, 11 bacterial colonies were selected for further analyses. Among the 11 strains, three strains were isolated from FW-B1, six strains were isolated from FW-B2, and two strains were isolated from FW-B3. The 16S rRNA sequencing suggested these 11 strains were *Alcaligenes faecalis*, *Myroides odoratimimus*, *Proteus terrae*, *Lysinibacillus fusiformis*, and *Paenochrobactrum gallinarii* (Table 2 and Figure 4). The three species isolated from FW-B1 were assigned to be *A. faecalis*, *M. odoratimimus*, and *P. gallinarii*. For the six strains isolated from FW-B2, one strain was *L. fusiformis*, and the other five strains were *P. terrae*. The two strains isolated from FW-B3 were identified to be *A. faecalis*.

**Figure 5.**
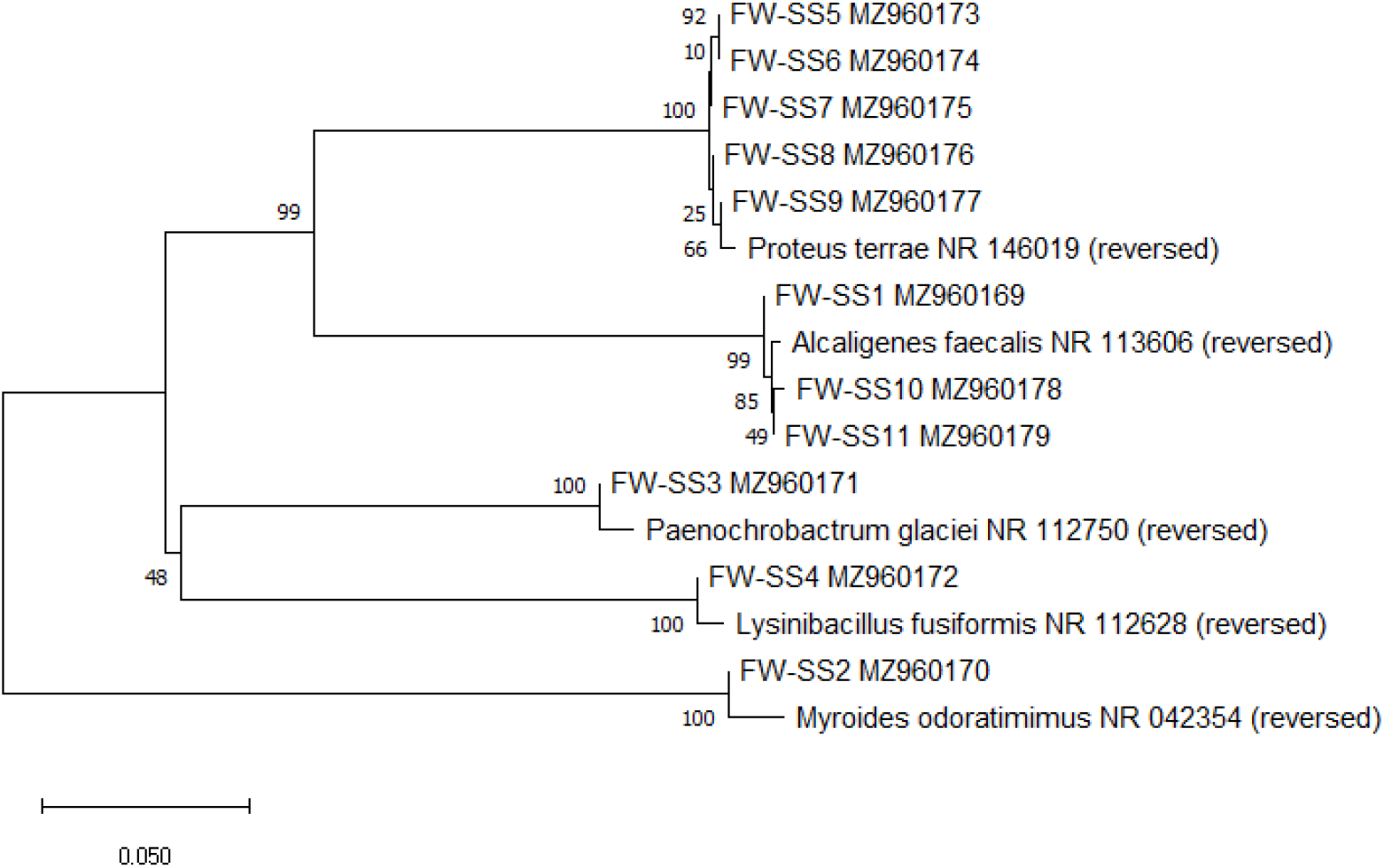
Phylogenetic tree of the 11 isolated strains.

**Table 2.** Isolated Bacteria from Initial Screening of Compost Samples

| Sample Number | GenBank Accession Number | Species Name (Accession Number) | Similarity |
| --- | --- | --- | --- |
| FW-SS1 | MZ960169 | *Alcaligenes faecalis* (NR_113606) | 99.55% |
| FW-SS2 | MZ960170 | *Myroides odoratimimus* (NR_042354) | 99.70% |
| FW-SS3 | MZ960171 | *Paenochrobactrum glaciei* (NR_112750) | 99.92% |
| FW-SS4 | MZ960172 | *Lysinibacillus fusiformis* (NR_112628) | 99.50% |
| FW-SS5 | MZ960173 | *Proteus terrae* (NR_146019) | 99.78% |
| FW-SS6 | MZ960174 | *Proteus terrae* (NR_146019) | 99.64% |
| FW-SS7 | MZ960175 | *Proteus terrae* (NR_146019) | 99.64% |
| FW-SS8 | MZ960176 | *Proteus terrae* (NR_146019) | 99.85% |
| FW-SS9 | MZ960177 | *Proteus terrae* (NR_146019) | 99.78% |
| FW-SS10 | MZ960178 | *Alcaligenes faecalis* (NR_113606) | 99.36% |
| FW-SS11 | MZ960179 | *Alcaligenes faecalis* (NR_113606) | 99.64% |

#### 3.3.2 Cultured Strains Functions

In order to screen functional strains for household food waste composting, we tried to isolate strains from a sewage wastewater treatment plant. A *Bacillus cereus* strain, OK058270, proved to have degradative ability for cellulose and lipid (Figure 3). We further measured the crude lipase activity of the *B. cereus* fermentation supernatant with tween 20, tween 60, and tween 80 as substrate, respectively. The average lipase activity for tween 20 was 164.40±17.28 U/mg; the average lipase activity for tween 60 was 180.68±23.40 U/mg; the average enzyme activity measured for tween 80 was 220.58±44.91 U/mg.

**Figure 6.**
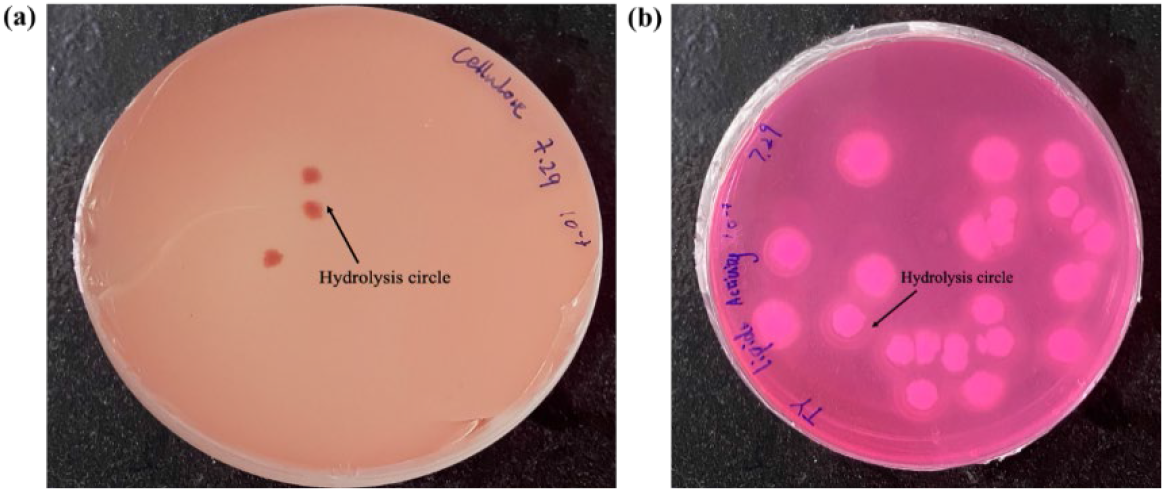
Clearance zones of *Bacillus cereus* strain on (a) cellulose and (b) lipid medium.

The isolated strain, *Bacillus cereus* OK058270, has been preserved by the China Centre for Type Culture Collection (CCTCC). CCTCC NO: M 20211679.

## 4. Discussion

We constructed three composts for efficient treatment of food waste at room temperature, and all the food waste were degraded in 81 days. During the 81-day composting process, the compost temperatures sustained at the room temperature, suggesting that the compost was at its mesophilic stage. Therefore, dominant strains identified in the compost samples would likely be the mesophilic species. Ammonia was present in the composting process, thus nitrifying species might be available in the compost microbiota. The microbiota of these three composts were revealed (Figure 2). Among the bacterial phyla of the FW-B1 to FW-B3 microbiota, *Firmicutes*, *Proteobacteria*, and *Bacteroidetes* were the three most dominant phyla. The identification of the three dominant bacterial phyla is consistent with other bacterial communities derived from food waste composts (Tran et al., 2020). These results suggest that the three phyla might be abundant in food waste composting. However, the phyla distribution of the three composts are different, showing that different food waste substrates might lead to different microbiota.

Moreover, the microbial difference between FW-B1 and FW-B3 might be due to the addition of commercial composting strains at the beginning of the composting process. This might be the reason *Lactobacillus* is the predominant genera in FW-B1, but not in FW-B3. It has also been reported that the EM commercial strains can remove odors caused by ammonia from a rubber factory. A large composition of *Lactobacillus* wass found in FW-B1, which might lead to less production of ammonia in FW-B1 (Figure 1b). Therefore, the addition of commercial strains could contribute to the odor elimination during composting, as well as the difference of the microbiota structures between FW-B1 and FW-B3.

Phylum *Firmicutes* has been widely distributed in the composting with various materials as substrate, including food waste and maize straw (Wei et al., 2018). Previous studies showed that *Firmicutes* are the most abundant phylum in the mesophilic stage of composting, but its abundance would gradually decline as the compost matures and enters the thermophilic stage (Chandna et al., 2013). This suggests that *Firmicutes* might be grown at the mesophilic stage of composting. During our composting process, the temperature of the composting box is dependent of the room temperature. This might account for the dominance of *Firmicutes* during the composting process.

At the genus level, the dominant genera is assigned to Firmicutes include *Lactobacillus*, *Peptoniphilus*, *Peptostreptococcus*, and *Clostridium. Lactobacillus* was the most dominant genus of all identified genera. *Lactobacillus* has the ability to increase composting efficiency, and thus would help the microbiota to degrade the waste (Li et al., 2020). Moreover, *Lactobacillus* strains isolated from composts can utilize carbohydrates, such as glucose and fructose, to produce lactic acid (Endo and Okada, 2007). All three composts were composed of a significant amount of carbohydrates, which could account for the dominance of *Lactobacillus. Lactobacillus* and *Peptoniphilus* were also shown to be associated with the production of malodorous gases (dimethyl disulfide) in food waste composts (Zhang et al., 2020). In real food waste treatment, decreasing their abundance might aid in the reduction of odors in composting, which would be particularly helpful in the household environment.

Some microbes assigned to *Proteobacteria* were involved in the ammonia-oxidizing process, and ammonia-oxidizing microbes have been identified in animal manure compost and other composts (Kowalchuk et al., 2020). During our composting process, pH tests suggest the presence of ammonia in the composts. This is consistent with high-level abundance of *Proteobacteria* in the compost bacterial community. *Proteobacteria* is also present in mesophilic environments (Fukuyama et al., 2019), and our composts had sustained at mesophilic temperatures. Dominant genera under phylum *Proteobacteria* include *Arcobacter*, *Proteus*, *Acinetobacter*, and *Comamonas. Arcobacter* is a gramnegative genus which can optimally grow at 30°C in microaerobic conditions (Banting and Salvat, 2017). *Arcobacter* species can be dominant at the early composting process, but it would disappear as the compost proceeds to the thermophilic stage (He et al., 2013). *Proteus* is another bacterial genus that is commonly found at the mesophilic stage in composting (Coker, 2019). *Acinetobacter* is known for its ability to accumulate antibiotic resistances, and previous studies showed that the dominance of *Acinetobacter* was lost after the composting process (Wang et al., 2015), suggesting that composting can help decrease potential pathogenic microbes. *Comamonas* is a strictly aerobic and mesophilic bacterial genus. The dominant proteobacteria genera identified from the composts were mostly mesophilic, and this might be due to the fact that our composting performs at the mesophilic condition.

The microbes assigned to phylum Bacteroidetes has been well known for their degradative capabilities of polymeric organic matter, and some Bacteroidetes were proved to be specialized in degrading polysaccharides and proteins (Thomas et al., 2011). The dominant genera of Bacteroidetes include *Myroides* and *Bacteroides*. Both of these two genera were dominant at the cooling phase of composting, when the composting is moving to maturity (Li et al., 2020; Li et al., 2019). The Bacteroides contributed up to 28% of the bacterial microbiota in composting of agricultural products (Li et al., 2020), suggesting that food waste compost microbiota were different with the agricultural substrate compost microbiota.

For the bacterial strains isolated from the compost samples, *A. faecalis*, *P. gallinarii*, and *P. terrae* belonged to phylum Proteobacteria, *M. odoratimimus* belonged to phylum Bacteroidetes, and *L. fusiformis* belonged to phylum *Fermicutes*. The isolated strains were all assigned to be from the three dominant phyla of the composts. *P. terrae* and *M. odoratimimus* were assigned to the dominant genera. However, strains of the most dominant genera, such as *Lactobacillus* and *Arcobacter*, were not isolated from the samples. This might be the a consequence of limitations to current isolation strategies.I In the future, culturomics can be applied to isolated dominant species in the food waste composts (Lagier et al. 2018).

In order to enhance compost performance, we tried to isolate functional strains from municipal wastewater treatment microbiota. A *B. cereus* strain with cellulase and lipase activities was isolated, and it has effective degradative capabilities of Tween 20, Tween 60, and Tween 80. In the future, *B. cereus* might be used as microbial agents to increase composting efficiency. We will add the culture to composts in future studies and evaluate the performance of the strain in a real composting environment.

## 5. Conclusion

In this study, the 16S rRNA gene fragment sequencing showed that *Firmicutes*, *Proteobacteria* and *Bacteroidetes* were the dominant phyla in the composting microbiota and *Lactobacillus*, *Arcobacter* and *Myroides* were the dominant genera. We also isolated five different microbial species from the microbiota. Moreover, in order to increase composting efficiency, we isolated one *Bacillus* strain with cellulase and lipase activities, which might be applied in future composting. We give a global insight into the composting microbiota, and future application of microbiome strategy would further increase composting efficiency. Characterization of the content of organic matter such as lipid, protein, cellulose, and starch in the compost material is needed in future studies to characterize the efficacy of the composting microbiota.

